# Platelet proteome analysis reveals an early hyperactive phenotype in SARS-CoV-2-infected humanized ACE2 mice

**DOI:** 10.1101/2021.08.19.457020

**Authors:** Saravanan Subramaniam, Ryan Matthew Hekman, Archana Jayaraman, Aoife Kateri O’Connell, Paige Montanaro, Benjamin Blum, Devin Kenney, Maria Ericsson, Katya Ravid, Nicholas A Crossland, Andrew Emili, Florian Douam, Markus Bosmann

**Affiliations:** Pulmonary Center, Department of Medicine, School of Medicine, Boston University MA, USA; Center for Network Systems Biology, Boston University, Boston, MA, USA; National Emerging Infectious Diseases Laboratories (NEIDL), Boston University, Boston, MA, USA; Department of Microbiology, School of Medicine, Boston University, Boston, MA, USA; Department of Pathology and Laboratory Medicine, School of Medicine, Boston University, Boston, MA, USA; Electron Microscopy Core Facility, Harvard Medical School, Boston, MA, USA; Department of Medicine, Boston University School of Medicine, Boston, MA, USA; Whitaker Cardiovascular Institute, Boston University School of Medicine, Boston, MA, USA; Center for Thrombosis and Hemostasis, University Medical Center of the Johannes Gutenberg-University, Mainz, Germany

## Abstract

Coronavirus disease-2019 (COVID-19) provokes a hypercoagulable state with increased incidence of thromboembolism and mortality. Platelets are major effectors of thrombosis and hemostasis. Suitable animal models are needed to better understand COVID-19-associated coagulopathy (CAC) and underlying platelet phenotypes. Here, we assessed K18-hACE2 mice undergoing a standardized SARS-CoV-2 infection protocol to study dynamic platelet responses via mass spectrometry-based proteomics. In total, we found significant changes in >1,200 proteins. Strikingly, protein alterations occurred rapidly by 2 days post-infection (dpi) and preceded outward clinical signs of severe disease. Pathway enrichment analysis of 2dpi platelet proteomes revealed that SARS-CoV-2 infection upregulated complement-coagulation networks (F2, F12, CFH, CD55/CD59), platelet activation-adhesion-degranulation proteins (PF4, SELP, PECAM1, HRG, PLG, vWF), and chemokines (CCL8, CXCL5, CXCL12). When mice started to lose weight at 4dpi, pattern recognition receptor signaling (RIG-I/MDA5, CASP8, MAPK3), and interferon pathways (IFIT1/IFIT3, STAT1) were predominant. Interestingly, SARS-CoV-2 spike protein in the lungs was observed by immunohistochemistry, but in platelets was undetected by proteomics. Similar to patients, K18-hACE2 mice during SARS-CoV-2 infection developed progressive lymphohistiocytic interstitial pneumonia with platelet aggregates in the lungs and kidneys. In conclusion, this model recapitulates activation of coagulation, complement, and interferon responses in circulating platelets, providing valuable insight into platelet pathology during COVID-19.

**Key Points:**

- SARS-CoV-2-infected humanized ACE2 mice recapitulate platelet reprogramming towards activation-degranulation-aggregation.
- Complement/coagulation pathways are dominant in platelets at 2 days post-infection (dpi), while interferon signaling is dominant at 4dpi.

## Introduction

Hospitalized COVID-19 patients frequently develop coagulation abnormalities with a high risk (∼10-40%) of thromboembolism.^1^ Elevated fibrin-degradation products are predictors of mortality from COVID-19,^2^ but therapeutic anticoagulation with heparin has achieved inconsistent survival benefits in clinical trials.^3,4^ Hence, there remains urgent need for molecular insight into COVID-19-associated coagulopathy (CAC), which shares some features with disseminated intravascular coagulopathy.^5^ During CAC, human platelets are activated and participate in procoagulatory-inflammatory responses to COVID-19.^6-8^ RNA-Seq has revealed what appears to be both direct and indirect effects (e.g. mediators, aberrant antibodies) of SARS-CoV-2 infection on the platelet transcriptome in patients.^9-12^

Mouse models are essential tools for drug development.^13^ While infected hACE2 mice present with microthrombosis,^14^ detailed platelet characterizations remain lacking. Here, we have phenotyped the platelet proteomes of SARS-CoV-2-infected humanized mice under standardized conditions and defined time points to provide an instrumental resource for future studies.

## Experimental design

Humanized ACE2 mice were infected intranasally with SARS-CoV-2. After 2 and 4 days, platelets were isolated from peripheral blood for quantitative proteomics, alongside histology and endpoints of clinical severity. (See supplementary materials for details.)

## Results and discussion

To evaluate K18-hACE2 mice for platelet alterations in CAC, we first characterized the time course of SARS-CoV-2 infection (1×10^6^ PFU). Body weight (Figure 1A), temperature (Figure 1B), clinical scores (supplemental Figure 1A), survival (Figure 1C), and viral loads (supplemental Figure 1B) started to deteriorate after 5dpi with high lethality at 7dpi.^14^

**Figure 1.**
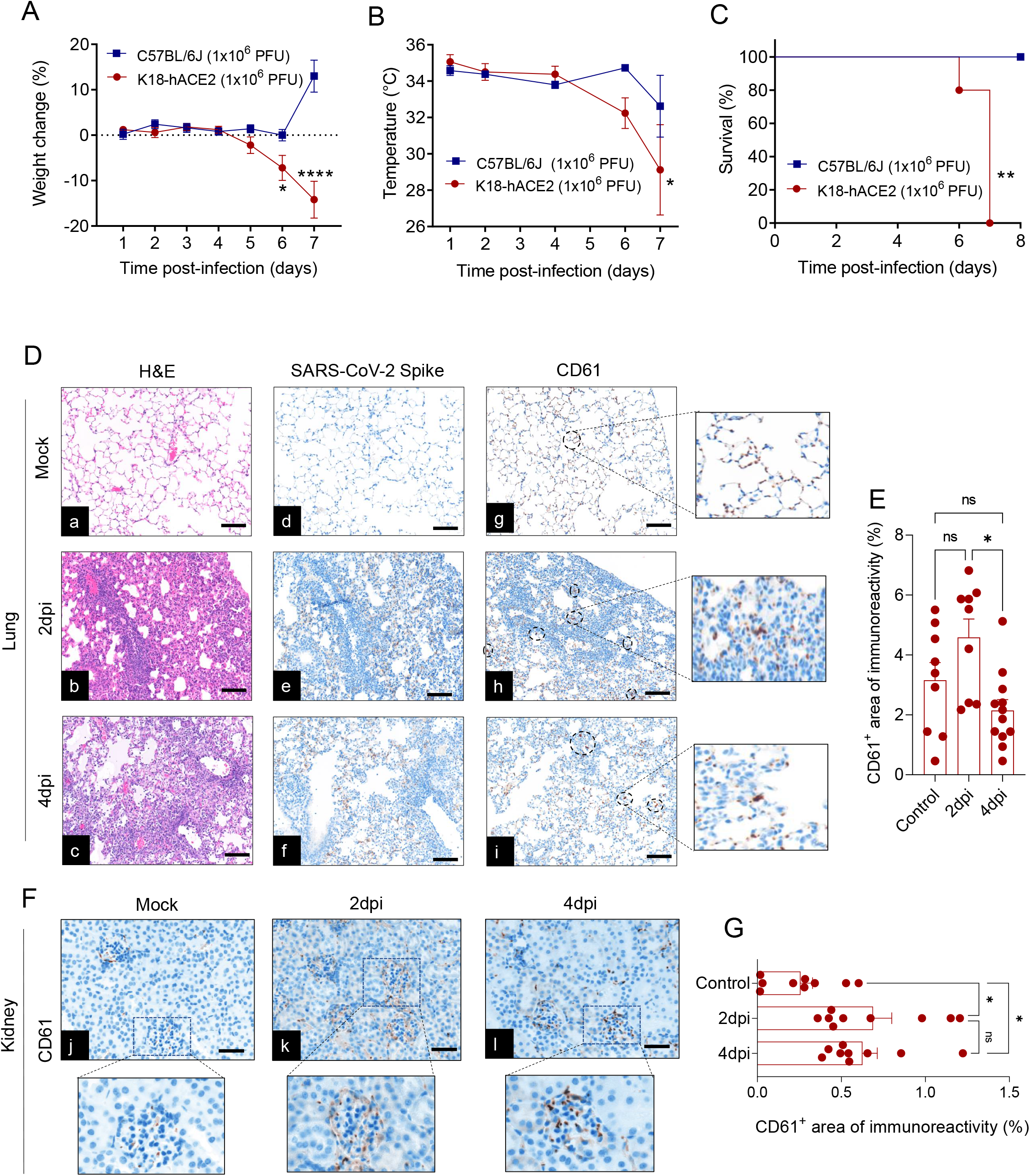
Microvascular platelet aggregates precede clinical decline of SARS-CoV-2-infected humanized ACE2 mice. K18-hACE2 mice and C57BL/6J mice were inoculated intranasally with 1×10^6^ plaque-forming units (PFU) or received saline (mock). (**A**) Body weight, (**B**) temperature, and (**C**) survival were monitored. (**D^a,b,c^**) Progressive interstitial pneumonia in K18-hACE2 mice at 2dpi and 4dpi with SARS-CoV-2, H&E staining. (**D^d,e,f^**) SARS-CoV-2 spike protein was detectable in the lungs of inoculated mice, immunohistochemistry. (**D^g,h,i^**) Lung capillaries contained increased CD61^+^ platelet aggregates in SARS-CoV-2-infected mice with no overt thrombosis. 200x, each image is representative of n=3 mice/group, scale bar=50 μm. (**E**) Whole-slide quantifications of platelet aggregates in lungs of SARS-CoV-2-infected mice. (**F^j-l^**) Renal capillaries contained increased platelet aggregates in SARS-CoV-2-inoculated mice with no overt thrombosis or edema. (**G**) Quantification of platelet aggregates in renal capillaries. Data are shown as the mean ± SEM, A-C: n=5 mice/group, D-G: n=3 mice/group, E, G: biological and technical triplicates; \**P < .05;* \*\**P < .01*; \*\*\*\**P < .0001; ns: not significant*.

As platelet-fibrin thrombi characterize multi-organ thrombosis of lethal COVID-19 in humans,^15^ we examined mouse lungs and kidneys at two time points (2dpi and 4dpi). Histologic findings included multifocal mild-to-moderate mononuclear cell infiltrates and low numbers of neutrophils in peribronchiolar, perivascular, interstitial, and alveolar spaces (Figure 1D^a-c^). SARS-CoV-2 spike protein was observed in the alveolar epithelium (Figure 1D^d-f^). Although there were no occlusive thrombi, we observed sporadic enhanced aggregations of CD61^+^ platelets in lung capillaries (Figure 1D^g-i^, 1E) and kidney interstitial/glomerular capillaries (Figure 1F^j-l^, 1G).^14^ Lung interstitial capillaries within areas of viral replication/assembly sporadically contained platelet aggregates (supplemental Figure 1C)^14^. These findings suggest platelet activation and aggregation is a feature of SARS-CoV-2 infection of K18-hACE2 mice already occurring before clinical decline.

Next, we analyzed platelet proteomes from SARS-CoV-2-infected K18-hACE2 mice at 2dpi/4dpi versus mock negative controls using mass spectrometry (Figure 2A). Principal component analysis illustrated clear differences between experimental groups (supplemental Figure 2A). In total, ∼4,600 platelet proteins were identified/quantified (supplemental File 1). SARS-CoV-2 induced significant changes in the abundance of many platelet proteins at both 2dpi (n=591↑/476↓; *P ≤ .05*; Figure 2B; supplemental File 2) and 4dpi (n=163↑/195↓; *P ≤ .05*; Figure 2C; supplemental File 3) compared to uninfected controls. Interestingly, we observed altered protein expression specific to 2dpi (n=376↑/470↓; *P ≤ .05*; Figure 2D, 2E; supplemental Figure 3) and 4dpi (n=95↑/42↓; *P ≤ .05*; Figure 2D, 2E; supplemental Figure 4) compared to mock platelets. In addition, many proteins were dysregulated at both 2dpi and 4dpi (n=100↑/121↓; *P ≤ .05*; Figure 2D, supplemental Figure 5). KEGG enrichment analysis revealed that complement and coagulation pathways were upregulated at 2dpi (Figure 2F), while intracellular infection pathways were upregulated at 4dpi (Figure 2G).

**Figure 2.**
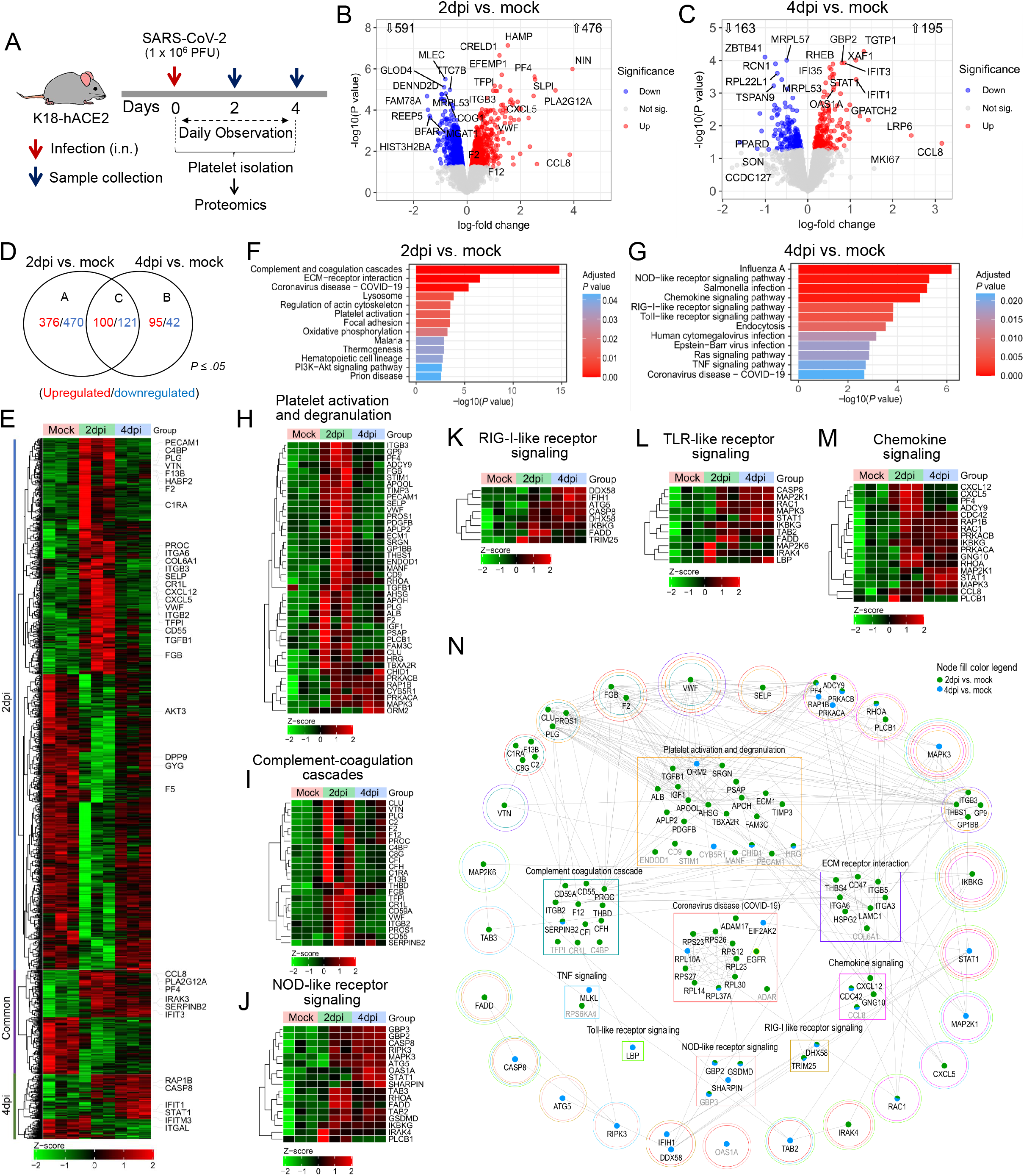
Distinct adaptations of platelet proteomes during SARS-CoV-2 infection. (**A**) K18-hACE2 mice were infected with SARS-CoV-2 (1×10^6^ PFU) and circulating platelets were collected at 2dpi and 4dpi for quantitative proteome analysis by mass spectrometry (biological triplicates). Volcano plots of differentially expressed proteins detected in (**B**) 2dpi versus mock and (**C**) 4dpi versus mock. Numbers of significantly (unadjusted *P ≤ .05*) up/downregulated proteins are shown. (**D**) Venn diagram and (**E**) heatmap of significantly regulated proteins (unadjusted *P ≤ .05*) in 2dpi versus mock, 4dpi versus mock or common for both time points. Raw normalized expression values (Z-score from −2/green to +2/red) and key proteins of interest are shown. (**F**) KEGG pathway enrichment analyses of significant (unadjusted *P ≤ .05*) upregulated proteins in 2dpi versus mock and (**G**) 4dpi versus mock. Pathways of interest with high statistical significance for enrichment (*FDR < .05*) are depicted. (**H**-**M**) Heatmaps of normalized expression (Z-score from −2/green to +2/red) of proteins (unadjusted *P ≤ .05*) comparing 2dpi versus mock and 4dpi versus mock platelet proteomics for the gene ontology pathways of (**H**) platelet activation and degranulation, (**I**) complement and coagulation pathways, (**J**) NOD-like receptor signaling, (**K**) RIG-I-like receptor signaling, (**L**) toll-like receptor signaling, and (**M**) chemokine signaling. (**N**) STRING-based protein-protein interaction network of differentially expressed proteins (unadjusted *P ≤ .05*) linked to platelet activation and degranulation, complement-coagulation cascades, chemokine signaling, RIG-I-like receptor signaling, toll-like receptor signaling, TNFα signaling, Coronavirus disease (COVID-19), ECM receptor interaction, or NOD-like receptor signaling. Protein nodes differentially expressed in 2dpi versus mock are colored green, 4dpi versus mock are in blue, and those altered in both time-points are colored with both green and blue. Proteins in square boxes were enriched only in one pathway/process while proteins within circles were enriched in more than one pathway/process. Circle colors represent the different pathways/processes. All data shown (B-N) are based on quantitative proteomics.

The analysis at 2dpi revealed a significant increase in SELP, PECAM-1, PLG, vWF ^16^ as well as thrombin expression (F2) (Figure 2H), supporting both the activation and degranulation of platelets during SARS-CoV-2 infection. Thrombin causes platelets to change shape, aggregate, and secrete prothrombotic granule contents. Elevated platelet thrombin might be due to circulating thrombin binding to platelet PAR3, which then activates platelet PAR4 during infection, although this needs further investigation. Similarly, an increase in coagulation factor F12 at 2dpi (Figure 2I) suggests plasma F12 may bind platelets to accelerate activation (contact pathway), and participate in CAC.^17^ Furthermore, reduced F5 at 2dpi (Figure 2E) could indicate F5 release to the plasma by activated platelets. On the other hand, we also observed increased anti-coagulation proteins (TFPI, PROC, PROS1, THBD) (Figure 2I), suggesting both pro-coagulant and anti-coagulant factors were prominently dysregulated during early SARS-CoV-2 infection.

Histidine-rich glycoprotein (HRG) is an abundant plasma protein with a multi-domain structure, allowing interaction with many ligands including phospholipids, plasminogen, fibrinogen, IgG antibodies, and heparan sulfate. HRG was significantly increased at 2dpi and 4dpi (Figure 2H). HRG exerts broad antiviral activities and might help to explain platelet-mediated antiviral responses during SARS-CoV-2 infection.^18^ In contrast, HRG may also have deleterious effects. We speculate that HRG-coated, activated platelets could adhere to inflamed endothelium via heparan sulfate, which would then block antithrombin-III binding and FXa inhibition, and could ultimately promote thromboembolism/microthrombi in arterioles and venules.

Platelet activation during thrombosis is closely associated with complement^19^ and contact system activation, antibody attack,^20^ and in turn inflammation. Interestingly, complement regulatory proteins (CD55, CD59, C1R, CFH, CFI) and membrane-attack complex component 8 (C8G) were significantly upregulated at 2dpi (Figure 2I). Similar to atypical hemolytic uremic syndrome,^21^ factor H (CFH) activity on platelets at 2dpi could be associated with platelet aggregation and activation. The significant increase in platelet CD55 and CD59 (GPI-anchored membrane proteins), and C1q-associated protease (C1R), support both complement system activation and platelet protection from complement-mediated destruction during SARS-CoV-2 infection.

Of note, significantly upregulated proteins at 4dpi were enriched in pathways typically engaged in the response to infection with viruses (Influenza A) and other intracellular pathogens (e.g., *Salmonella*). These included NOD-like receptor signaling (CASP8, GBP3, GBP2, MAPK3) (Figure 2J), and Toll-like and RIG-I-like receptor signaling pathways (DDX58/RIG-I, IFIH1/MDA5, DHX58, IRAK4, RAC1, MAPK3) (Figure 2K, 2L). In line with this, we observed dominant interferon-induced proteins (IFIT1, IFITM3) and STAT1 in platelets at 4dpi, suggesting a platelet immune response (Figure 2E). In addition to these pathways, certain chemokines (PF4, CCL8) (Figure 2M) and TNFα signaling components (RIPK3, CASP8) (supplemental Figure 2B) were strongly upregulated at both 2dpi and 4dpi compared to uninfected controls. PF4/CXCL4 is of particular interest as the most abundant platelet kinocidin. It is released from α-granules of activated platelets^22^. Importantly, platelets from COVID-19 patients have been shown to release δ/α-granule cargo into the blood.^11^ PF4 has also been implicated in vaccine-induced thrombotic responses.^23^ In addition, CXCL12, CXCL5, and CCL8 were likewise elevated (Figure 2M). These chemokines recruit monocytes/macrophages to infection sites and promote platelet-leukocyte aggregates.^24^

A protein-protein interaction network of significantly upregulated proteins at 2dpi and 4dpi was retrieved from the STRING database, which revealed modules with multiple sources of biological crosstalk (Figure 2N). For instance, vWF has been reported to play a critical role in COVID-19 pulmonary microvascular occlusion.^25^ In our interactome, we noted high interconnectivity between vWF, PLG, PF4, ITGB3 and MAPK3, as well as between PF4, CXCL12 and CXCL5. Dysregulation of vWF, F2, and PF4 may serve as central nodes in enhancing virus-induced cytopathology similar to RSV,^26^ by interacting with proteins governing platelet activation and degranulation, complement activation, coagulation, ECM-receptor interaction, and other key pathways dysregulated in COVID-19. Of note, proteins upregulated at 2dpi were more interconnected compared to 4dpi (Figure 2N).

Within our K18-hACE2 mouse platelet samples, SARS-CoV-2 proteins, ACE2, and TMPRSS2 were not detectable by mass spectrometry (supplemental File 1). In human platelets, SARS-CoV-2 uptake seems infrequent, independent of ACE2/TMPRSS2, and can induce cell death pathways.^6,9-11,27^ Apoptotic proteins were also upregulated in mouse platelets (supplemental Fig. 2C). In conclusion, our study identified platelet proteome signatures dependent on SARS-CoV-2 infection timing, and validated K18-hACE2 mice as a suitable model for future studies gaging the direct impacts of specific platelet components on COVID-19 progression.

## Author contributions

M.B., F.D., and S.S. conceptualized the overall study. K.R. and A.M. contributed to testable hypotheses. F.D., A.O., D.K., and S.S. performed experiments. R.Y. performed mass spec analyses of SARS-CoV-2-infected platelet samples. N.A.C. and P.M. performed histology and immunohistochemistry analyses. R.Y., A.J., and S.S. performed proteome data analyses. M.E. performed transmission electron microscopy of SARS-CoV-2-infected platelet samples. S.S., M.B., N.A.C., A.E., and F.D. interpreted data. S.S., M.B., and K.R. wrote and/or revised the manuscript. All authors read and approved the manuscript.

## Acknowledgments

This work was supported by the National Institutes of Health (1R01HL141513, 1R01HL139641, 1R01AI153613, 1UL1TR001430 to M.B.; 1R21ES032882, 1K22AI144050 to F.D.), a Boston University Start-up fund (to F.D.), a Peter Paul Career Development Professorship (to F.D.), and utilized a Ventana Discovery Ultra autostainer that was purchased with funding from a National Institutes of Health grant: S10 OD026983 (PAR-18-600). We thank the Evans Center for Interdisciplinary Biomedical Research at Boston University School of Medicine for their support of the Affinity Research Collaborative on ‘Respiratory Viruses: A Focus on COVID-19’. We thank Ulrich Walter and Lucien Peter Garo for reading the manuscript and Bernhard Lämmle for advice. The authors are responsible for the content of this publication.

## Disclosure

M.B. and F.D. are funded by ARCA Biopharma for another project on COVID-19.

## Figure Legends

**Figure S1.**
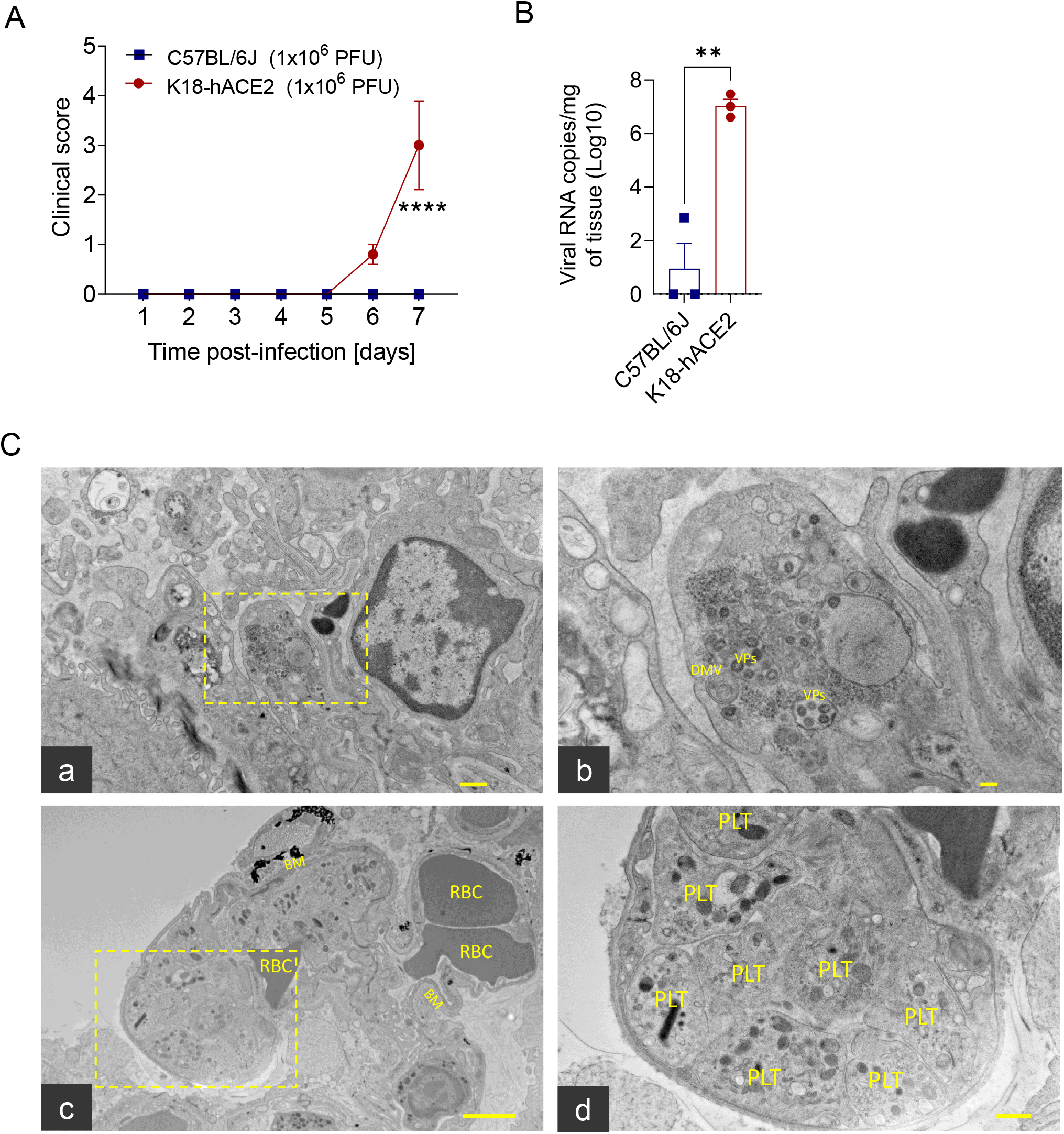
Clinical scores, infection load, and ultrastructural findings of lung cross sections from SARS-CoV-2 infected K18-hACE2 mice. K18-hACE2 mice and C57BL/6J controls were inoculated with SARS-CoV-2 (1×10^6^ PFU, i.n.) and monitored for the indicated time. (**A**) Clinical scores, n=5/group. (**B**) Viral load in lungs at 6dpi, n=3/group. (**C^a-b^**) Viral assembly within an alveolar type I pneumocyte as evidenced by presence of double membrane-bound vesicles (DMVs) that routinely contain virus particles (VPs), transmission electron microscopy. (**C^c-d^**) Interstitial capillaries adjacent to areas of viral assembly occasionally containing aggregates of platelets at 6dpi. RBC: red blood cells; BM: basement membrane; Scale bars in frames C^a-d^: a=500 nm; b=100 nm; c=2 μm; d=500 nm. Data are shown as mean ± SEM; \*\**P* < *.01*. \*\*\*\**P < .0001*.

**Figure S2.**
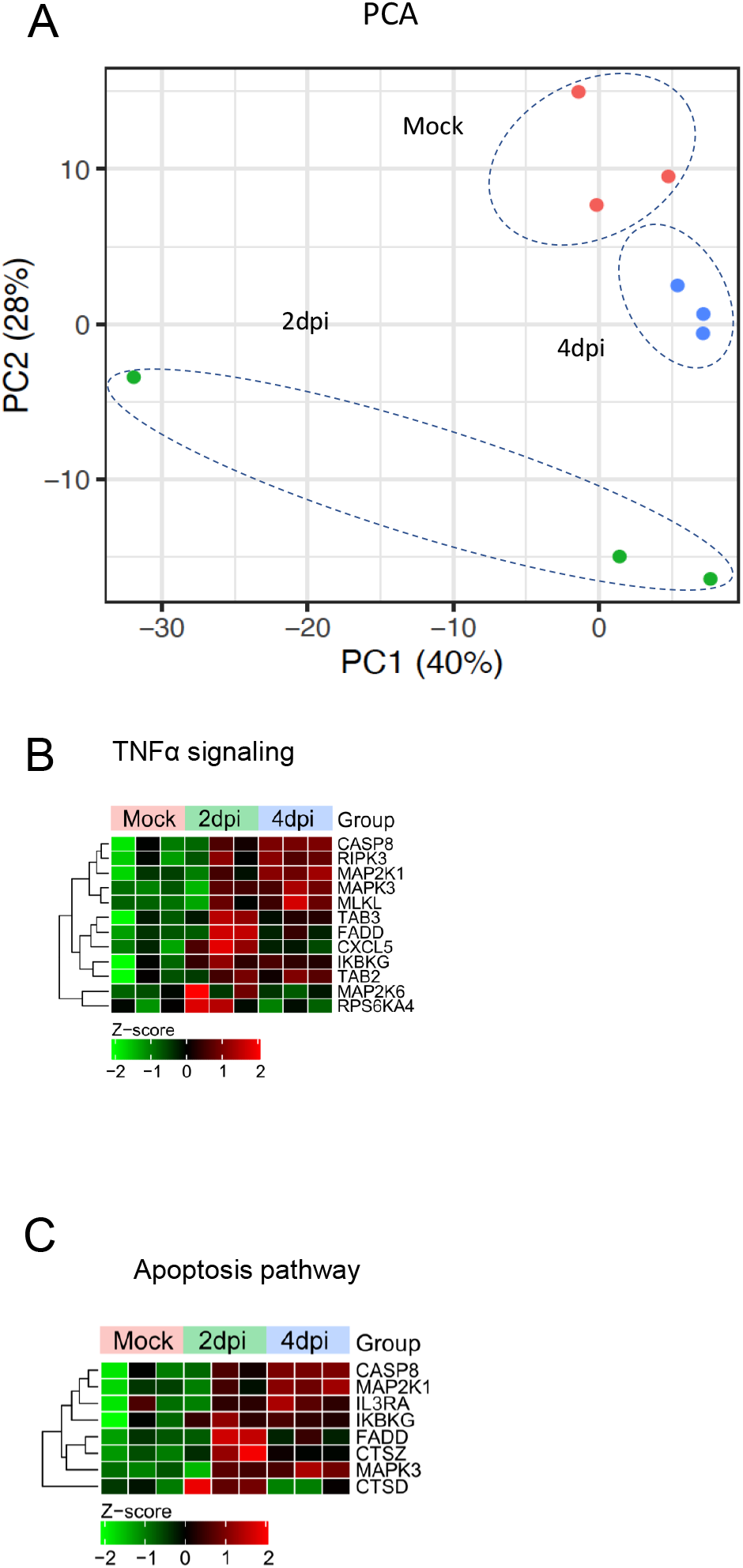
Platelet proteome analysis. (**A**) Principal component analysis (PCA) plot showing proteomic variance (%) in mock, 2dpi, and 4dpi quantitative proteomics datasets along PC1 and PC2 in K18-hACE2 mice (SARS-CoV-2, 1×10^6^ PFU i.n.; biological triplicates). (**B**-**C**) Heatmaps of normalized expression (Z-score from −2/green to +2/red) of proteins (unadjusted *P ≤ .05*) in (**B**) TNF-α signaling and (**C**) the apoptosis pathway, comparing 2dpi versus mock and 4dpi versus mock proteomics data.

**Figure S3.**
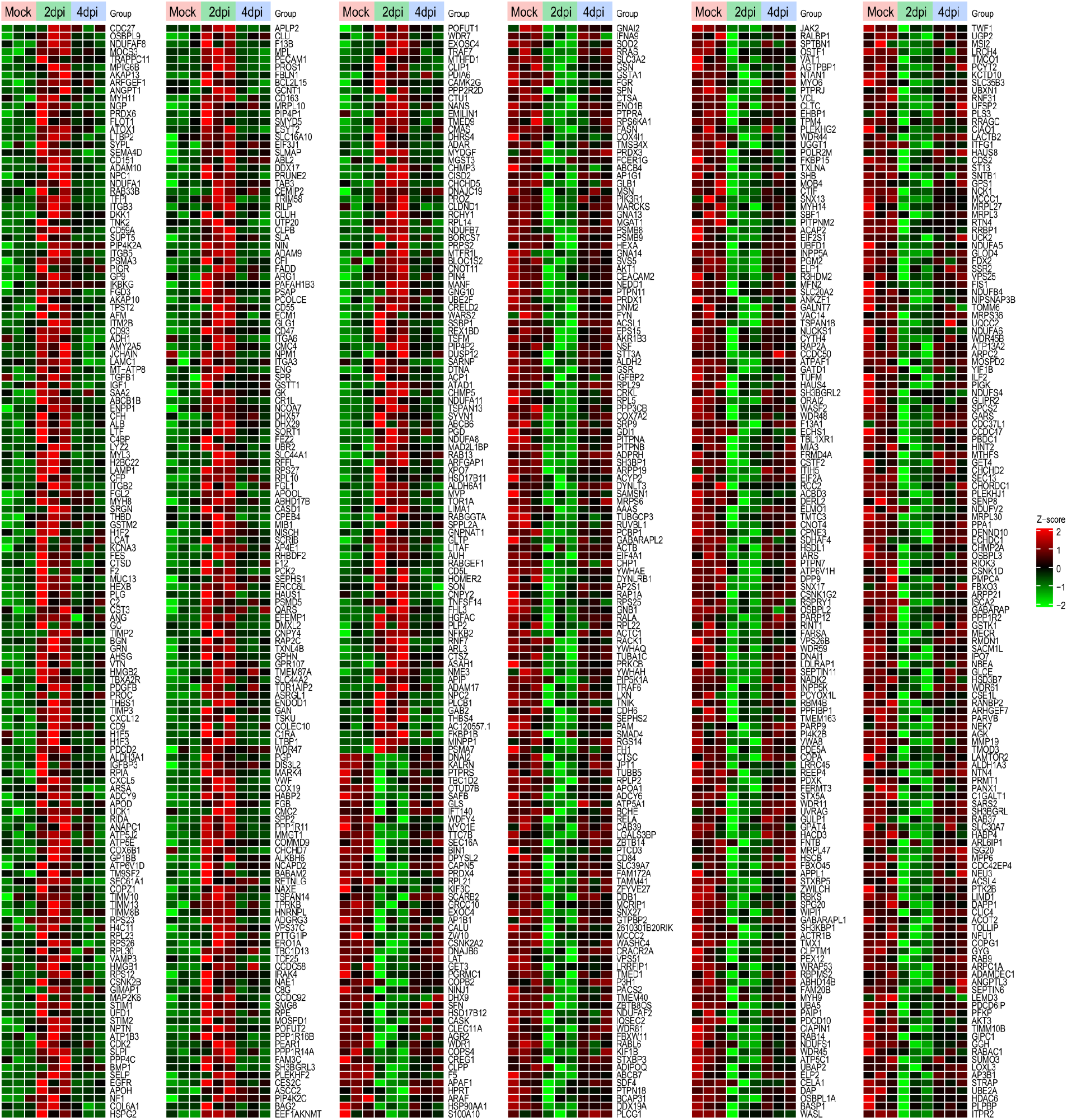
Significantly regulated proteins specific to 2dpi. Heatmaps of normalized expression (Z-score from −2/green to +2/red) of significantly upregulated and downregulated proteins in the platelet proteomics dataset (unadjusted *P ≤ .05; venn section A from figure 2D*) in 2dpi K18-hACE2 mice compared to uninfected K18-hACE2 controls (mock); biological triplicates for all groups. The data for 4dpi versus mock were not significant for all proteins shown.

**Figure S4.**
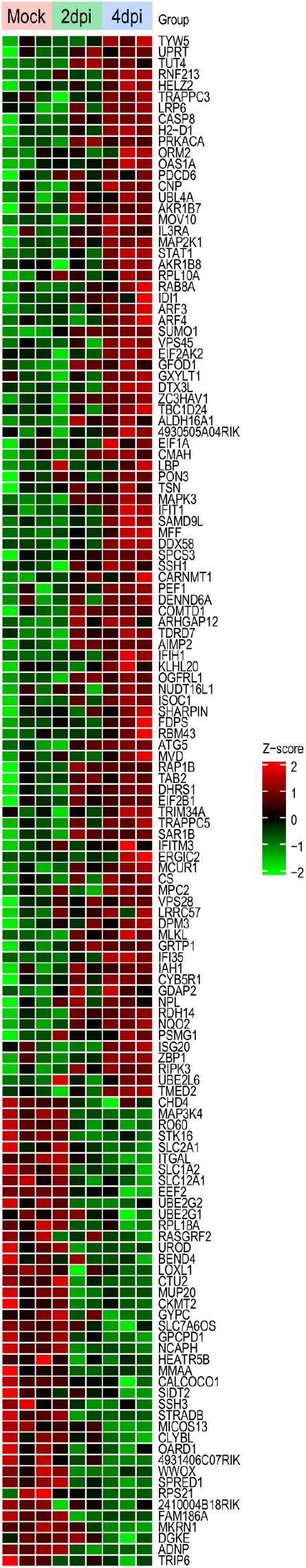
Significantly regulated proteins specific to 4dpi. Heatmaps of normalized expression (Z-score from −2/green to +2/red) of significantly upregulated and downregulated proteins in the platelet proteomics dataset (unadjusted *P ≤ .05; venn section B from figure 2D*) in 4dpi SARS-CoV-2 infected K18-hACE2 mice compared to uninfected K18-hACE2 controls (mock); biological triplicates for all groups. The data for 2dpi versus mock were not significant for all proteins shown.

**Figure S5.**
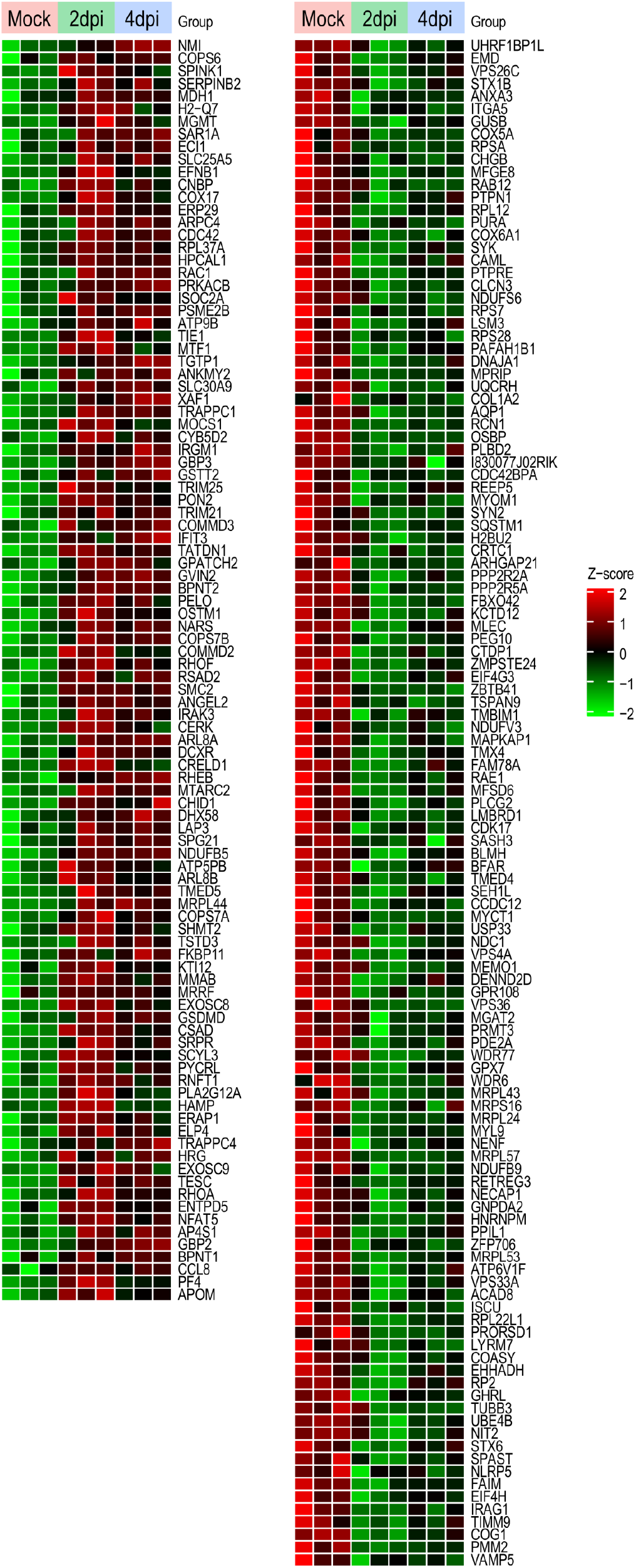
Significantly regulated proteins common to 2dpi and 4dpi. Heatmaps of normalized expression (Z-score from −2/green to +2/red) of common significantly upregulated and downregulated platelet proteins (unadjusted *P ≤ .05; venn section C from figure 2D*) in 2dpi versus mock and 4dpi versus mock proteomics datasets (biological triplicates).

## Supplemental materials

### Methods

#### Ethics statement

All experimental procedures with animals were approved by the Boston University Biomedical Research, Institutional Biosafety Committee and Institutional Animal Care and Use Committee.

#### SARS-CoV-2 and propagation

2019-nCoV/USA-WA1/2020 isolate (NCBI accession number: MN985325) of SARS-CoV-2 was obtained from the Centers for Disease Control and Prevention (Atlanta, GA) and BEI Resources (Manassas, VA). African green monkey kidney Vero E6 cells (ATCC^®^ CRL-1586^™^, American Type Culture Collection, Manassas, VA) were used to propagate the viral isolate *in vitro*. Two and three days after infection, the culture medium was collected, filtered through a 0.2 µm filter, and ultra-centrifuged (Beckman Coulter Optima L-100k; SW32 Ti rotor) for 2 h at 25,000 rpm (80,000*g*) over a 20% sucrose cushion (Sigma-Aldrich, St. Louis, MO). Pellets were resuspended overnight at 4 °C in 500 μl of 1X PBS. The next day, concentrated virions were aliquoted and stored at −80 °C. The work with SARS-CoV-2 was performed in a biosafety level-3 laboratory by personnel equipped with powered air-purifying respirators at the National Emerging Infectious Diseases Laboratories (NEIDL), Boston University.

#### Virus titration

The titer of our viral stock was determined by plaque assay. In brief, Vero E6 cells were seeded into a 12-well plate at a density of 2.5×10^5^ cells per well. The next day, the cells were infected with serial 10-fold dilutions of the virus stock for 1 h at 37 °C. After virus adsorption, 1 ml of overlay media (2X DMEM supplemented with 4% FBS and mixed at a 1:1 ratio with 1.2% Avicel (DuPont; RC-581) was added in all the wells. Three day later, the overlay medium was removed, the cell monolayer was washed with 1X PBS and fixed with 4% paraformaldehyde for 30 min at room temperature. Next, cells were washed with 1X PBS and stained with 0.1% crystal violet in 10% ethanol/water for 1 h at room temperature. The stained wells were rinsed with tap water and the numbers of plaques were counted. The titer of our passage (P)-2 virus stock was 4×10^8^ plaque formatting units (PFU) per ml.

#### Mice

C57BL/6J mice and heterozygous humanized ACE2 (K18-hACE2) mice (strain: 2B6.Cg-Tg(K18-ACE2)2Prlmn/J) were obtained from the Jackson Laboratory (Bar Harbor, ME). Animals were maintained in Tecniplast green line individually ventilated cages (Tecniplast, Buguggiate, Italy). Mice were maintained on a 12:12 light cycle at 30-70% humidity and provided *ad libitum* water and standard chow diets (LabDiet, St. Louis, MO).

#### SARS-CoV-2 infection

14 week old C57BL6J and K18-hACE2 transgenic female mice were intranasally inoculated with 1×10^6^ PFU of SARS-CoV-2 in 50 μl of sterile 1X PBS and sham mice were inoculated with 50 μl of sterile 1X PBS. Inoculations were performed under 1-3% isoflurane anesthesia. Survival studies were performed for 7 days post infection (dpi). Mice were euthanized early if they reached euthanasia criteria, or if the experiment included predetermined time points for sample collection (2dpi, 4dpi).

#### Clinical monitoring

The mice of the 7-day survival studies were intraperitoneally implanted with a radio-frequency identification (RFID) temperature-monitoring microchip (Unified Information Devices, Lake Villa, IL, USA) 48-72 hours prior to inoculation. An IACUC-approved clinical scoring system was used to monitor disease progression and establish humane endpoints ^1^. The evaluated categories were body weight, general appearance, responsiveness, respiration, and neurological signs. Clinical signs and body temperature were recorded once daily for the full duration of the studies.

#### Viral RNA isolation

Lung and kidney tissues were collected from mice and stored in 600 μl of RNAlater (Sigma-Aldrich; #R0901500ML) and stored at −80 °C. For RNA isolation, 20-30 mg of tissue were placed into 2 ml tubes with 600 μl of RLT buffer with 1% β-mercaptoethanol and 5 mm stainless steel beads (Qiagen, Valencia, CA; #69989). Tissues were then dissociated using a Qiagen TissueLyser II (Qiagen) with the following cycle parameters: 20 cycles/s for 2 min, 1 min wait, 20 cycles/s for 2 min. Samples were centrifuged at 13,000 rpm (17,000*g*) for 10 minutes and supernatants were transferred to new 1.5 ml tubes. Viral RNA isolation was performed using a Qiagen RNeasy Plus Mini Kit (Qiagen; #74134), according to the manufacturer’s instructions.

#### SARS-CoV-2 E-specific reverse transcription quantitative polymerase chain reaction (RT-qPCR)

Viral RNA was quantitated using single-step RT-quantitative real-time PCR (Quanta qScript One-Step RT-qPCR Kit, QuantaBio, Beverly, MA; VWR; #76047-082) with primers and TaqMan® probes targeting the SARS-CoV-2 E gene as previously described ^1^. Briefly, a 20 μL reaction mixture containing 10 μL of Quanta qScriptTM XLT One-Step RT-qPCR ToughMix, 0.5 μM Primer E_Sarbeco_F1 (ACAGGTACGTTAATAGTTAATAGCGT), 0.5 μM Primer E_Sarbeco_R2 (ATATTGCAGCAGTACGCACACA), 0.25 μM Probe E_Sarbeco_P1 (FAM-ACACTAGCCATCCTTACTGCGCTTCG-BHQ1), and 2 μL of template RNA was prepared. RT-qPCR was performed using an Applied Biosystems QuantStudio 3 (ThermoFisher Scientific) and the following cycling conditions: reverse transcription for 10 minutes at 55 °C, an activation step at 94 °C for 3 min followed by 45 cycles of denaturation at 94 °C for 15 seconds and combined annealing/extension at 58 °C for 30 seconds. Ct values were determined using QuantStudio Design and Analysis software V1.5.1 (ThermoFisher Scientific). For absolute quantitation of viral RNA, a 389 bp fragment from the SARS-CoV-2 E gene was cloned onto pIDTBlue plasmid under an SP6 promoter using NEB PCR cloning kit (New England Biosciences, Ipswich, MA). The cloned fragment was then *in vitro* transcribed (mMessage mMachine SP6 transcription kit; ThermoFisher) to generate an RT-qPCR standard.

#### Platelet isolation and sample preparation

Mice were anesthetized by placement in a chamber connected to a 1-3% isoflurane vaporizer. The unconscious mice were injected intraperitoneally with a lethal dose of ketamine/xylazine for euthanasia. Citrated whole blood was immediately collected by heart puncture and supplemented with 1 ml of modified Tyrode’s buffer (134 mM NaCl, 12 mM NaHCO_3_, 2.9 mM KCl, 0.34 mM Na_2_HPO_4_, 1 mM MgCl_2_, 10 mM HEPES, pH 6.5). Platelet-rich plasma was prepared by centrifugation at 100*g* for 10 minutes at room temperature. After two washes with modified Tyrode’s buffer pH 6.5, cells were lysed in a lysis buffer comprising 6M GuHCl, 100 mM Tris pH 8.0, 40 mM chloroacetamide and 10 mM TCEP, supplemented with Complete Mini protease inhibitor (Roche; #11836170001) and phosphatase inhibitor cocktails (Roche; #04906837001). This isolation method consistently provided ≥95% pure platelets and less than 0.05% contaminating leukocytes as confirmed by flow cytometry (data not shown). For virus inactivation, lysates were heated to 100 °C for 15 min, snap frozen on dry ice and stored at −80 °C. Total proteins (∼100 µg/sample for mass spectrometry) were quantified using Bradford’s assay and normalized prior to digestion with trypsin (Pierce, ThermoFisher Scientific) at 1:50 ratio (enzyme to protein, w/w) overnight at 37 °C. Digestion was quenched with trifluoracetic acid (pH 3) and the peptides were desalted using reverse-phase Sep-Pak C18 columns (Waters Corporation; WAT054955) with a wash buffer of 0.1% TFA and elution buffer of 60% acetonitrile. The desalted peptides were then quantified with a Quantitative Colorimetric Peptide Assay (Pierce). Desalted peptides (20 μg) were labeled with Tandem Mass Tags (TMT) using TMTPro-16plex isobaric tags (ThermoFisher Scientific; #A44520) as per manufacturer’s instructions. Triplicate samples corresponding to each time point were labelled separately.

TMT-labeled peptides were fractionated via basic reversed-phase chromatography on the Agilent 1200 series HPLC instrument equipped with the XBridge Peptide BEH C18 column (130A°, 3.5 mm, 4.6 mm X 250 mm, Waters Corporation). Prior to loading peptides, the C18 column was washed with 100% methanol and equilibrated with Buffer A (0.1% ammonium hydroxide and 2% acetonitrile). Peptides were injected via the autosampler and eluted from the column using a gradient of mobile phase A (2% ACN, 0.1% NH4OH) to mobile phase B (98% ACN, 0.1% NH4OH) over 48 min at a flow-rate of 0.4 ml/minute. The 48 fractions collected were orthogonally concatenated into 12 pooled fractions. Five percent of each fraction was aliquoted and saved for global proteomic profiling. All fractions were dried in SpeedVac Vacuum Concentrator (Thermo Fisher Scientific).

#### Mass spectrometry analysis

Multiplexed peptide fractions from each time point were resuspended in mobile phase A solvent (2% acetonitrile and 0.1% formic acid) for analysis on an Exploris 480 mass spectrometer equipped with FAIMS (Thermo Fisher Scientific). The mass spectrometer was interfaced to the Easy nanoLC1200 HPLC system (Thermo Fisher Scientific). Briefly, the peptides were first loaded onto a reverse-phase nanotrap column in mobile phase A (Acclaim PepMap100 C18, 3 µm, 100 A°; 75 µm i.d. x 32 cm, Thermo Fisher Scientific) and then separated over an EASY-Spray capillary column, (ES803A; Thermo Fisher Scientific) using a gradient (6% to 19% over 58 min, then 19% to 36% over 34 min) of mobile phase B (0.1% formic acid, 80% acetonitrile) at a flow rate of 250 nl/min. The mass spectrometer was operated in positive ion mode with a capillary temperature of 275 °C and a spray voltage of 2500 V. All data was acquired with the mass spectrometer operating in data-dependent acquisition (DDA) mode, with FAIMS cycling through one of three compensation voltages (−50V, −57V, −64V) at each full scan. Precursor scans were acquired at a resolution of 120,000 FWHM with a maximum injection time of 120 ms in the Orbitrap analyzer. The following 0.8s were dedicated to fragmenting the topmost abundant ions at the same FAIMS compensation voltage, with charge states between 2 and 5, via HCD (NCE 33%) before analysis at a resolution of 45,000 FWHM with a maximum injection time of 60 ms.

#### Analysis of raw mass spectrometry data

All acquired MS/MS spectra were simultaneously searched against the complete SwissProt mouse proteome (downloaded on 2020-10-20) and the Uniprot SARS-CoV-2 proteome (downloaded on 2020-05-03) using MaxQuant (Version 1.6.7.0), which integrates the Andromeda search engine. TMT reporter ion quantification was performed using MaxQuant with default settings. Briefly, enzyme specificity was set to trypsin and up to two missed cleavages were allowed. Cysteine carbamidomethylation was specified as fixed modification whereas oxidation of methionine and N-terminal protein acetylation were set as variable modifications. Precursor ions were searched with a maximum mass deviation of 4.5 ppm and fragment ions with a maximum mass deviation of 20 ppm. Peptide and protein identifications were filtered at 1% FDR using the target-decoy database search strategy ^2^. Proteins that could not be differentiated based on MS/MS spectra alone were grouped to protein groups (default MaxQuant settings). The MaxQuant output files designated and ‘‘protein groups’’ were used for data normalization and other statistical analysis using in-house generated scripts in the R environment.

#### Transmission electron microscopy

Tissue samples were fixed for 72 hours in a mixture of 2.5% Glutaraldehyde and 2% formaldehyde in 0.1 M sodium cacodylate buffer (pH 7.4). Samples were then washed in 0.1 M cacodylate buffer and postfixed with 1% osmium tetroxide (OsO4) / 1.5% potassium ferrocyanide (KFeCN6) for 1 hour at room temperature. After washes in water and 50 mM maleate buffer pH 5.15 (MB), the samples were incubated in 1% uranyl acetate in MB for 1 h, washed in MB and water, and dehydrated in grades of alcohol (10 min each; 50%, 70%, 90%, 2×10min 100%). The tissue samples were then put in propylene oxide for 1 h and infiltrated overnight in a 1:1 mixture of propylene oxide and TAAB Epon (TAAB Laboratories Equipment Ltd, Aldermaston, Berks, RG7 8NA, England). The following day the samples were embedded in fresh TAAB Epon and polymerized at 60°C for 48 h. Semi-thin (0.5 μm) and ultrathin sections (50-80 nm) were cut on a Reichert Ultracut-S microtome (Leica, Charleston, SC, USA). Semi-thin sections were picked up on glass slides and stained with Toluidine blue for examination at the light microscope level to find affected areas in the tissue. Ultrathin sections from those areas were picked up onto formvar/carbon coated copper grids, stained with 0.2% lead citrate and examined in a JEOL 1200EX transmission electron microscope (JOEL, Akishima, Tokyo, Japan). Images were recorded with an AMT 2k CCD camera (AMT Imaging Woburn, MA 01801).

#### Histology and immunohistochemistry

Animals were anesthetized with 1-3% isoflurane and euthanized with an intraperitoneal overdose of ketamine and xylazine before vital organ removal and fixation of tissues. Lungs were insufflated with ∼1.5 ml of 1% low melting point agarose (Sigma-Aldrich) diluted in 1X PBS using a 18-gauge catheter placed into the trachea. Tissues were inactivated in 10% neutral buffered formalin at a 20:1 fixative to tissue ratio for a minimum of 72 h before removal from BSL-3 in accordance with an approved institutional standard operating procedure. Tissues were subsequently processed and embedded in paraffin following standard histological procedures. Sections (5 µm) were obtained and stained with hematoxylin and eosin (H&E). Immunohistochemistry (IHC) was performed using a Ventana BenchMark Discovery Ultra autostainer (Roche Diagnostics, Indianapolis, IN) using ChromoMap DAB kit (Roche Diagnostics). PBS mock animals served as biologic negative controls for SARS-CoV-2 S-protein, while the absence of immunoreactivity was confirmed by the occlusion of primary antibodies or use of an isotype control specific to human tissues. The following antibodies were used: Mouse-SARS-CoV-2 spike protein (clone: E7U6O; Cell Signaling Technology, USA); rabbit-CD61 (clone: ARC0460; Invitrogen, USA). Quantification of both H&E and IHC stainings was performed in biological and technical triplicates.

#### Whole slide image analysis

Slides were imaged on the HALO™ image analysis platform using a Mantra 2.0^™^ quantitative pathology workstation and unmixed with InForm software version 2.4.8 (Akoya Biosciences, Marlborough, MA, USA). For each mouse, 3 images of lung tissue and 3 images of kidney tissue were obtained. Slides were manually annotated to adjust visualization thresholds in order to fine-tune visibility of markers within each sample. Due to inconsistencies in lung insufflation, the Classifier module within HALO was used to discern lung stroma and glass within tissue images. Quantitative outputs of CD61^+^ staining were obtained using the Area Quantification (AQ) module, which reports total area of immunoreactivity of a specified parameter within a region of interest. For kidney samples, the entire image was analyzed. For lung samples, only the regions classified as lung stroma were analyzed, glass space was excluded. Values were expressed as a percentage of total tissue area analyzed. AQ was performed to determine percentage of CD61^+^ immunoreactivity in kidneys and lungs of the studied mice.

#### Statistical analysis

Statistical analysis was performed using GraphPad Prism 9 software (GraphPad Software, LLC) by one-way ANOVA (multiple groups) and Bonferroni multiple-comparisons test, or student-t test (2 groups). Survival rates were analyzed by Log-rank (Mantel-Cox) test. *P* values *< .05* were considered statistically significant. Bioinformatics analysis (pathway enrichment analysis) and data visualization of significant differentially expressed proteins (*P* value [rounded to two decimal places] *< .05*) were performed using the R framework (v4.0.2) in RStudio Server (v1.3.1073). Entrez and Ensembl IDs were mapped using biomaRt (v2.44.4) and only proteins with mapped IDs were used for visualization and downstream analysis. Volcano plots were constructed using ggplot2 (v3.3.3). Venn diagrams were constructed using VennDiagram (v1.6.20) and VennDetail (v1.4.0). Heatmaps were generated using ComplexHeatmap (v2.4.3) package with unsupervised clustering, and the dendrograms were reordered for a more meaningful visualization using dendsort (v0.3.3) and dendextend (v1.14.0) packages. Gene Ontology (GO), including biological process (BP), molecular function (MF) and cellular component (CC), was analyzed using enrichR (v3.0) and gProfiler2 (v0.2.0). Pathway analyses based on Kyoto Encyclopedia for Genes and Genomes (KEGG) and Reactome were performed using clusterProfiler (v3.16.1) and ReactomePA (v1.32.0), respectively. False discovery rate (FDR) was controlled using Benjamini– Hochberg method. All functional enrichment analyses were performed based on pathways specific to *Mus musculus* using gene symbols. Plots were generated for GO and pathway enrichment specific to significantly upregulated proteins. To understand interactions between proteins in nine key processes and pathways that were enriched in either 2dpi or 4dpi data sets, an interactome was constructed and visualized using the stringApp (v1.6.0) in Cytoscape (v3.8.2). The network was specific to *Mus musculus* and included only predicted interactions with the highest combined confidence score (≥0.900). We mapped the logFold change values for these proteins onto the interactome.

#### Data sharing statement

The proteomics data presented in this manuscript and raw data files will be deposited in the ProteomeXchange Consortium (http://www.proteomexchange.org) upon the acceptance of the manuscript. All other data are available from the corresponding author upon reasonable request.

## Notes

### Competing Interest Statement

The authors have declared no competing interest.

